# Phage infection and sub-lethal antibiotic exposure mediate *Enterococcus faecalis* type VII secretion system dependent inhibition of bystander bacteria

**DOI:** 10.1101/2020.05.11.077669

**Authors:** Anushila Chatterjee, Julia L. E. Willett, Gary M. Dunny, Breck A. Duerkop

**Affiliations:** Department of Immunology and Microbiology, University of Colorado School of Medicine, Aurora, CO, USA, 80045.; Department of Microbiology and Immunology, University of Minnesota Medical School, Minneapolis, MN, USA, 55455.

**Keywords:** bacteriophages, *Enterococcus*, antibiotic resistance, phage–bacteria interactions, bacterial secretion systems, type VII secretion, contact-dependent antagonism

## Abstract

Bacteriophages (phages) are being considered as alternative therapeutics for the treatment of multidrug resistant bacterial infections. Considering phages have narrow host-ranges, it is generally accepted that therapeutic phages will have a marginal impact on non-target bacteria. We have discovered that lytic phage infection induces transcription of type VIIb secretion system (T7SS) genes in the pathobiont *Enterococcus faecalis*. Membrane damage during phage infection induces T7SS gene expression resulting in cell contact dependent antagonism of different Gram positive bystander bacteria. Deletion of *essB*, a T7SS structural component, abrogates phage-mediated killing of bystanders. A predicted immunity gene confers protection against T7SS mediated inhibition, and disruption of its upstream LXG toxin gene rescues growth of *E. faecalis* and *Staphylococcus aureus* bystanders. Phage induction of T7SS gene expression and bystander inhibition requires IreK, a serine/threonine kinase, and OG1RF_11099, a predicted GntR-family transcription factor. Additionally, sub-lethal doses of membrane targeting and DNA damaging antibiotics activated T7SS expression independent of phage infection, triggering T7SS antibacterial activity against bystander bacteria. Our findings highlight how phage infection and antibiotic exposure of a target bacterium can affect non-target bystander bacteria and implies that therapies beyond antibiotics, such as phage therapy, could impose collateral damage to polymicrobial communities.

**Author Summary:** Renewed interest in phages as alternative therapeutics to combat multi-drug resistant bacterial infections, highlights the importance of understanding the consequences of phage-bacteria interactions in the context of microbial communities. Although it is well established that phages are highly specific for their host bacterium, there is no clear consensus on whether or not phage infection (and thus phage therapy) would impose collateral damage to non-target bacteria in polymicrobial communities. Here we provide direct evidence of how phage infection of a clinically relevant pathogen triggers an intrinsic type VII secretion system (T7SS) antibacterial response that consequently restricts the growth of neighboring bacterial cells that are not susceptible to phage infection. Phage induction of T7SS activity is a stress response and in addition to phages, T7SS antagonism can be induced using sub-inhibitory concentrations of antibiotics that facilitate membrane or DNA damage. Together these data show that a bacterial pathogen responds to diverse stressors to induce T7SS activity which manifests through the antagonism of neighboring non-kin bystander bacterial cells.

## Introduction

Enterococci constitute a minor component of the healthy human microbiota [1]. Enterococci, including *Enterococcus faecalis*, are also nosocomial pathogens that cause a variety of diseases, including sepsis, endocarditis, surgical-site, urinary tract and mixed bacterial infections [2, 3]. Over recent decades, enterococci have acquired extensive antibiotic resistance traits, including resistance to “last-resort” antibiotics such as vancomycin, daptomycin, and linezolid [4–8]. Following antibiotic therapy, multi-drug resistant (MDR) enterococci can outgrow to become a dominant member of the intestinal microbiota, resulting in intestinal barrier invasion and blood stream infection [7, 9]. The ongoing evolution of MDR enterococci in healthcare settings [4–6, 10, 11] and their ability to transmit antibiotic resistance among diverse bacteria [9, 12–15], emphasize the immediate need for novel therapeutic approaches to control enterococcal infections.

Viruses that infect and kill bacteria (bacteriophages or phages) are receiving attention for their use as antibacterial agents [16]. Recent studies have demonstrated the efficacy of anti-enterococcal phages in murine models of bacteremia [17–19] and the administration of phages to reduce *E. faecalis* burden in the intestine gives rise to phage resistant isolates that are sensitized to antibiotics [20]. Considering phages are highly specific for their target bacterium, coupled with the self-limiting nature of their host-dependent replication, this suggests that unlike antibiotics which have broad off-target antimicrobial activity, phages should have nominal impact on bacteria outside of their intended target strain [21–23]. However, our understanding of how phages interact with bacteria and the bacterial response to phage infection is limited.

While studying the transcriptional response of phage infected *E. faecalis* cells, we discovered that phage infection induces the expression of genes involved in the biosynthesis of a type VIIb secretion system (T7SS) [24]. Firmicutes, including the enterococci, harbor diverse T7SS genes encoding transmembrane and cytoplasmic proteins involved in the secretion of protein substrates [25], and T7SSs promote antagonism of non-kin bacterial cells through production of antibacterial effectors and/or toxins [26, 27]. The antibacterial activity of T7SSs from staphylococci and streptococci are well characterized [25] but T7SS-mediated antibacterial antagonism has not been described for enterococci. The environmental cues and regulatory pathways that govern T7SS expression and activity are poorly understood, although recent studies indicate that exposure to serum and membrane stresses triggered by pulmonary surfactants, fatty acids and phage infection stimulate T7SS gene expression [24, 28–31]. This motivated us to determine if phage induced T7SS gene expression in *E. faecalis* results in the inhibition of non-kin bacterial cells that are not phage targets (bystanders). We discovered that phage infected *E. faecalis* produces potent T7SS antibacterial activity against bystander bacteria. Expression of a T7SS antitoxin (immunity factor) gene in bystander cells and mutation of the LXG domain containing gene located immediately upstream of this immunity factor confer protection against phage mediated T7SS inhibition. We also investigated the potential impact of antimicrobials directed against bacterial physiological processes that are also targeted by phages, including cell wall, cell membrane and DNA damaging agents, on enterococcal T7SS. Sub-lethal challenge with specific antibiotics enhances T7SS gene expression resulting in T7SS dependent interspecies antagonism. Additionally, we discovered that membrane stress during phage infection induces transcription of T7SS genes via a non-canonical IreK signaling pathway. To our knowledge, the enterococcal T7SS is the first example of secretion system induction during phage infection. These data shed light on how phage infection of a cognate bacterial host can influence polymicrobial interactions and raises the possibility that phages may impose unintended compositional shifts among bystander bacteria in the microbiota during phage therapy.

## Results

### Phage mediated induction of *E. faecalis* T7SS leads to interspecies antagonism

A hallmark feature of phage therapy is that phages often have a narrow host range, hence they do not influence the growth of non-susceptible bacteria occupying the same niche [22]. We discovered that infection of *E. faecalis* OG1RF by phage VPE25 induces the expression of T7SS genes [24]. The *E. faecalis* OG1RF T7SS locus is absent in the commonly studied vancomycin-resistant strain V583, despite conservation of flanking genes (Fig. 1A) [32, 33]. The OG1RF T7SS is found downstream of conserved tRNA-Tyr and tRNA-Gln genes, which could facilitate recombination or integration of new DNA [34], but no known recombination or integration sites were identified on the 3’ end of this locus. Homologs of the *E. faecalis* T7SS gene *esxA* are found throughout three of the four *Enterococcus* species groups [35], including *Enterococcus faecium*, suggesting a wide distribution of T7SS loci in enterococci (Fig. S1). In addition to EsxA, OG1RF encodes the core T7SS structural components EsaA, EssB, and EssC, which are predicted to localize to the membrane, and EsaB, a small predicted cytoplasmic protein (Fig. 1A) [36]. OG1RF_11102 encodes an additional putative membrane protein, although it does not share sequence homology with staphylococcal or streptococcal EssA. We were unable to identify an EssA homolog in OG1RF using sequence-based homology searches, suggesting that the enterococcal T7SS machinery may differ from previously described T7SS found in other Gram-positive bacteria. *In silico* analyses predict that the *E. faecalis* T7SS locus encodes multiple WXG100 family effectors and LXG family polymorphic toxins [27, 37]. We hypothesized that induction of T7SS genes during phage infection and consequently the heightened production of T7SS substrates would indirectly influence the growth of non-kin phage-resistant bacterial cells.

**Figure 1.**
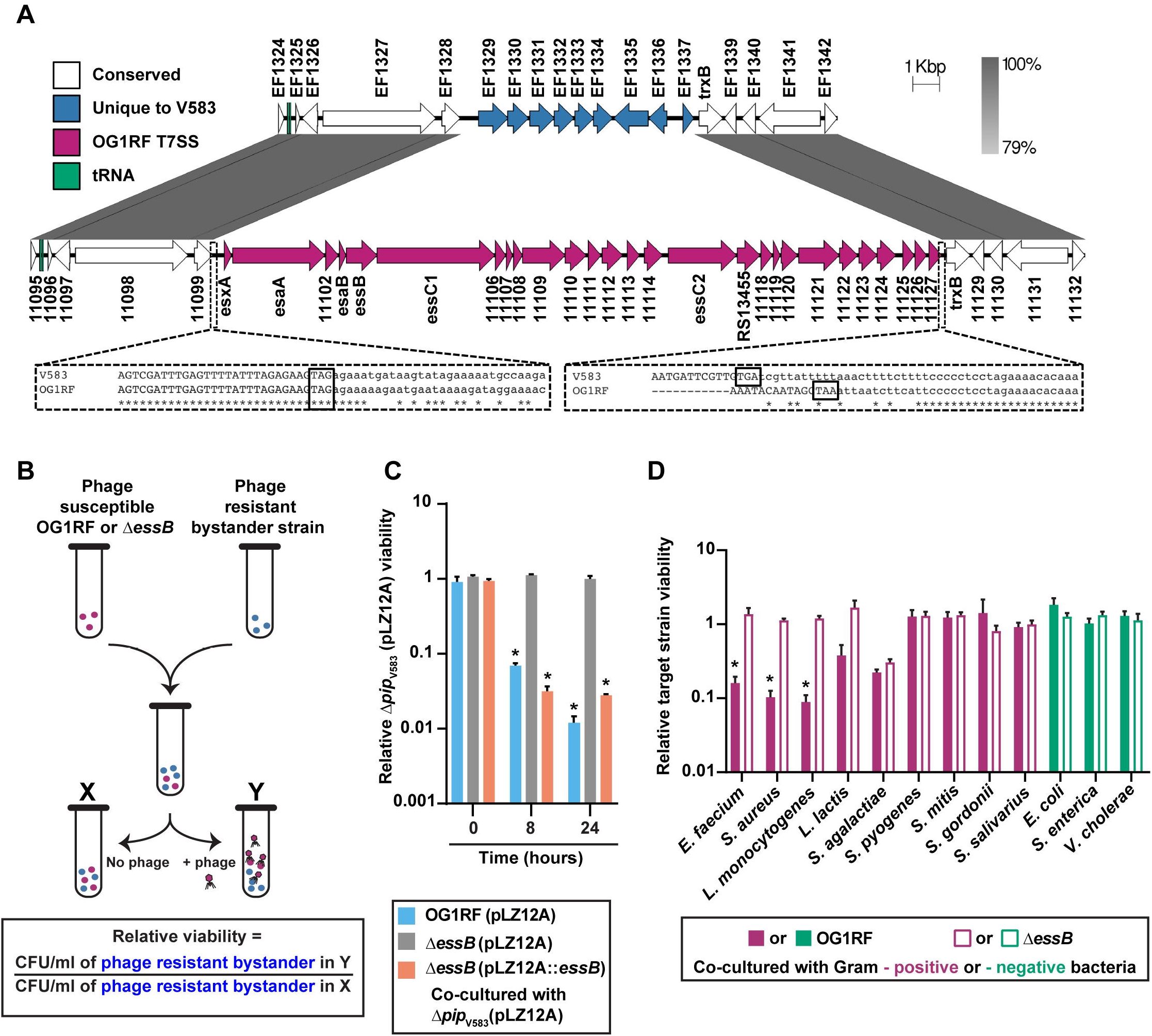
Phage mediated inhibition of bystander bacteria is dependent on enterococcal T7SS. **(A)** Diagram showing the location of T7SS genes in *E. faecalis* OG1RF (NC_017316.1) compared to *E. faecalis* V583 (NC_004668.1). Sequences were obtained from NCBI, and homology comparisons were rendered in EasyFig. Nucleotide alignments generated by Clustal Omega are enlarged for clarity (dashed lines). Stop codons of genes EF1328/OG1RF_11099 and EF1337/OG1RF_11127 are boxed. **(B)** Schematic representation of the co-culture assay used to assess the viability of bystander bacteria during phage induced T7SS activity of wild type *E. faecalis* OG1RF and Δ*essB*. Relative viability of bystander strains is calculated by measuring the ratio of bystander cfus in the phage infected culture compared to the bystander cfus from an uninfected control culture. **(C)** The relative abundance of viable bystander bacterium *E. faecalis* Δ*pip*_V583_. Complementation of the *E. faecalis* Δ*essB* mutant, Δ*essB* (pLZ12A::*essB*), restores T7SS dependent bystander inhibition. Δ*essB* (pLZ12A) is the empty vector control. **(D)** T7SS inhibition of other bacterial species in the presence and absence of phage infected *E. faecalis* OG1RF or Δ*essB*. Data represent three biological replicates. Error bars indicate standard deviation. \**P* < 0.00001 by unpaired Student’s t-test.

To investigate if T7SS factors produced during phage infection of *E. faecalis* OG1RF interferes with the growth of phage-resistant bystander bacteria, we generated a strain with an in-frame deletion in the T7SS gene *essB*, encoding a transmembrane protein involved in the transport of T7SS substrates [38]. We chose to inactivate *essB* as opposed to the more commonly investigated secretion promoting ATPase *essC* [27, 38], because *E. faecalis* OG1RF harbours two *essC* genes in its T7SS locus that may have functional redundancy (Fig. 1A). The *essB* mutant is equally susceptible to phage VPE25 infection compared to wild type *E. faecalis* OG1RF (Fig. S2A). We performed co-culture experiments where phage susceptible wild type *E. faecalis* OG1RF or Δ*essB* were mixed with a phage resistant bystander, a strain of *E. faecalis* V583 deficient in the production of the VPE25 receptor (Δ*pip_V583_*) [39], at a ratio of 1:1 in the absence and presence of phage VPE25 (multiplicity of infection [MOI] = 0.01) (Fig. 1B). VPE25 infected *E. faecalis* OG1RF and the Δ*essB* mutant with similar efficiency and caused a 1000-fold reduction in the viable cell count over a period of 24 hours relative to the starting cell count (Fig. S2B). Since sequence-based homology searches did not retrieve any homologs of potential antitoxins from the *E. faecalis* OG1RF T7SS locus in *E. faecalis* V583 genome, this strain likely lacks immunity to toxins encoded in this locus. The viability of *E. faecalis* Δ*pip*_V583_, was reduced nearly 100-fold when co-cultured with *E. faecalis* OG1RF in the presence of phage VPE25 (Fig. 1C, S2C). However, growth inhibition of *E. faecalis* Δ*pip*_V583_ was abrogated during co-culture with phage infected *E. faecalis* Δ*essB* and phage induced T7SS antagonism of *E. faecalis* Δ*essB* could be restored by complementation (Fig. 1C, S2C), indicating that inhibition of phage resistant *E. faecalis* Δ*pip*_V583_ by OG1RF is T7SS dependent.

T7SS encoded antibacterial toxins secreted by Gram positive bacteria influence intra- and interspecies antagonism [26, 27]. While a nuclease and a membrane depolarizing toxin produced by *Staphylococcus aureus* target closely related *S. aureus* strains [26, 40], *Streptococcus intermedius* exhibits T7SS dependent antagonism against a wide-array of Gram positive bacteria [27]. To determine the target range of *E. faecalis* OG1RF T7SS antibacterial activity, we measured the viability of a panel of VPE25 insensitive Gram positive and Gram negative bacteria in our co-culture assay (Fig. 1B). Growth inhibition of the distantly related bacterial species *E. faecium* and Gram positive bacteria of diverse genera, including *S. aureus* and *Listeria monocytogenes*, occurred following co-culture with phage infected wild type *E. faecalis* OG1RF but not the Δ*essB* mutant (Fig. 1D). Fitness of *Lactococcus lactis*, a lactic acid bacterium like *E. faecalis*, was modestly reduced during co-culture with phage infected *E. faecalis* OG1RF, although these data were not statistically significant. In contrast, Gram positive pathogenic and commensal streptococci were unaffected (Fig. 1D). Similarly, phage induced T7SS activity did not inhibit any Gram negative bacteria tested (Fig. 1D). Collectively, these results show that phage predation of *E. faecalis* promotes T7SS inhibition of select bystander bacteria.

### Molecular basis of *E. faecalis* phage–triggered T7SS antagonism

Our data demonstrate that induction of *E. faecalis* OG1RF T7SS genes during phage infection hinder the growth of select non-kin bacterial species. Antibacterial toxins deployed by Gram negative bacteria via type V and VI secretion and Gram positive T7SS require physical contact between cells to achieve antagonism [26, 27, 41, 42]. Therefore, we investigated if growth inhibition of bystander bacteria is contingent upon direct interaction with phage infected *E. faecalis* using a trans-well assay [27]. We added unfiltered supernatants from wild type *E. faecalis* OG1RF and Δ*essB* mutant cultures grown for 24 hrs in the presence and absence of phage VPE25 (MOI = 0.01) to the top of a trans well and deposited phage resistant *E. faecalis* Δ*pip*_V583_ in the bottom of the trans well. The 0.4 μm membrane filter that separates the two wells is permeable to proteins and solutes but prevents bacterial translocation. Supernatant from phage infected wild type *E. faecalis* OG1RF did not inhibit *E. faecalis* Δ*pip*_V583_ (Fig. S3A) indicating that T7SS mediated growth interference relies on cell to cell contact. To exclude the possibility that T7SS substrates might adhere to the 0.4 μm membrane filter in the trans-well assay, we administered both filtered and unfiltered culture supernatants directly to *E. faecalis* Δ*pip*_V583_ cells (5×10^5^ CFU/well) at a ratio of 1:10 (supernatant to bystander cells) and monitored growth over a period of 10 hours. Growth kinetics of *E. faecalis* Δ*pip*_V583_ remained similar irrespective of the presence or absence of conditioned supernatant from wild type *E. faecalis* OG1RF or Δ*essB* mutant cultures (Fig. S3B – S3C), further supporting the requirement of contact-dependent engagement of phage mediated T7SS inhibition.

We discovered that *E. faecalis* OG1RF inhibits proliferation of non-kin bacterial cells through increased expression of T7SS genes in response to phage infection, but the toxic effectors were unknown. LXG domain containing toxins are widespread in bacteria with a diverse range of predicted antibacterial activities [43, 44]. The OG1RF T7SS locus encodes two LXG-domain proteins, OG1RF_11109 and OG1RF_11121 (Fig. 2A). Both LXG domains were found using Pfam, but we were unable to identify predicted function or activity for either protein using sequence homology searches or structural modeling.

**Figure 2.**
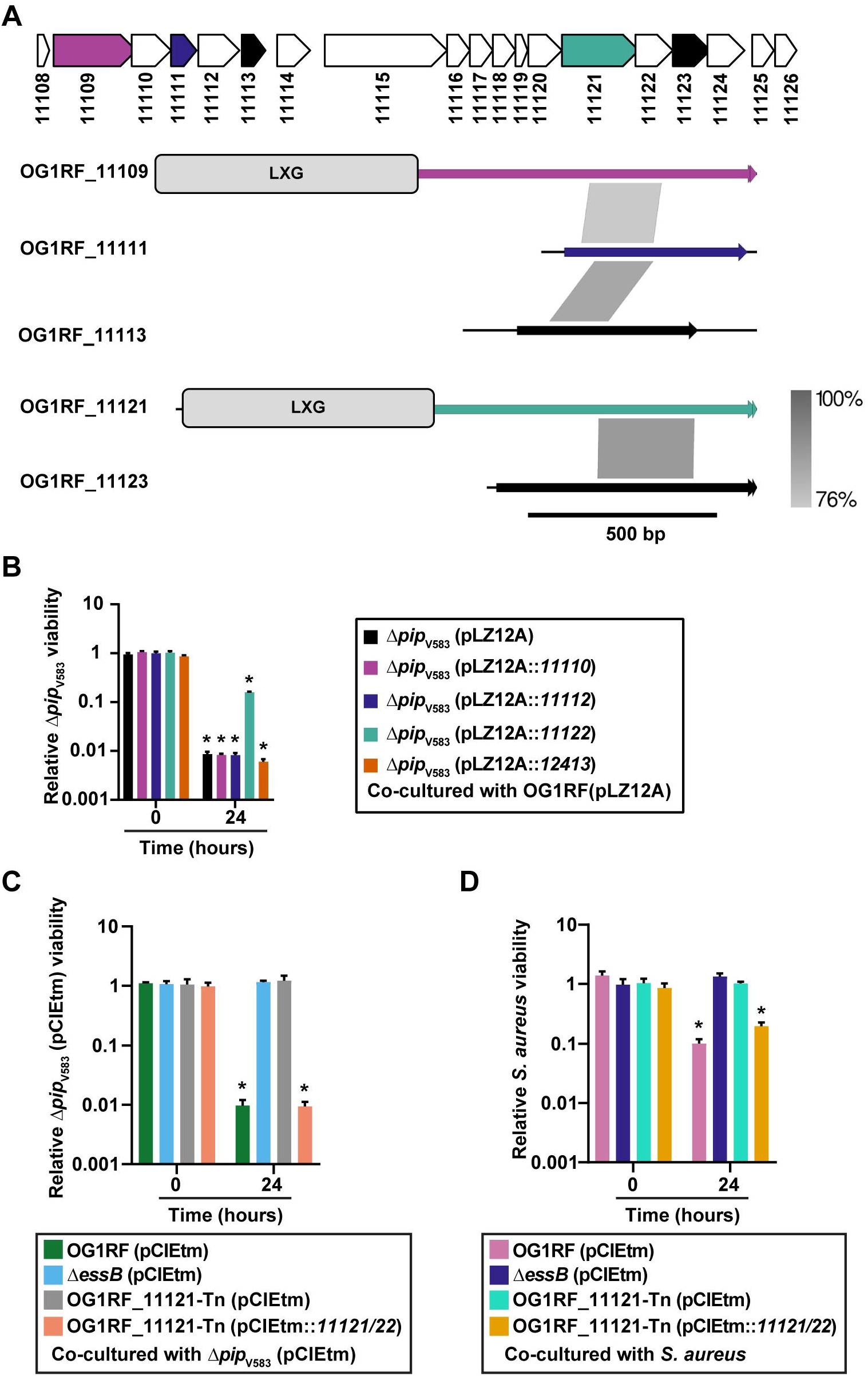
Identification of *E. faecalis* T7SS toxin and immunity proteins that dictate bystander growth inhibition. **(A)** Putative toxin-encoding genes in the OG1RF T7SS locus. LXG domains in OG1RF_11109 and OG1RF_11121 were identified using KEGG and ExPASy PROSITE. Putative orphan toxins were identified by homology to OG1RF_11109 or OG1RF_11121. Gray lines between diagrams indicate the regions and degree of nucleotide conservation between genes. Homology diagrams were rendered in EasyFig. Gene colors for OG1RF_11109, OG1RF_11111, and OG1RF_11121 match the color scheme in panel **(B)**. OG1RF_11113 and OG1RF_11123 are shaded black to indicate that their corresponding immunity genes were not tested in panel **(B)**. **(B)** *E. faecalis* OG1RF T7SS mediated growth inhibition of phage resistant *E. faecalis* Δ*pip*_V583_ during infection is alleviated by expressing OG1RF_11122 in *E. faecalis* Δ*pip*_V583_ but not in the presence of pLZ12A empty vector, or expressing OG1RF_11110, OG1RF_11112, or OG1RF_12413. **(C – D)** Disruption of OG1RF_11121 by a transposon insertion rescues growth of phage resistant *E. faecalis* Δ*pip*_V583_ **(C)** and *S. aureus* **(D)** strains during co-culture. Complementation of OG1RF_11121-Tn restores bystander intoxication. Data represent three biological replicates. Error bars indicate standard deviation. \**P* < 0.0001 by unpaired Student’s t-test.

Bacterial polymorphic toxin systems can encode additional toxin fragments and cognate immunity genes, known as “orphan” toxin/immunity modules, downstream of full-length secreted effectors [45, 46]. Orphan toxins lack the N-terminal domains required for secretion or delivery, although they can encode small regions of homology that could facilitate recombination with full-length toxin genes [47]. Therefore, we sought to identify putative orphan toxins in OG1RF. We aligned the nucleotide sequences of OG1RF_11109 and OG1RF_11121 with downstream genes in the T7SS locus and looked for regions of similarity that might signify orphan toxins. Although the 3’ ends of OG1RF_11111 and OG1RF_11113 did not have homology to either OG1RF_11109 or OG1RF_11121, the 5’ ends of OG1RF_11111 and OG1RF_11113 had >75% nucleotide homology to a portion of OG1RF_11109 (Fig. 2A, regions of homology indicated by gray shading). Similarly, OG1RF_11123 had sequence homology to OG1RF_11121 (Fig. 2A). We searched Pfam and ExPasy for annotated domains but were unable to identify any in OG1RF_11111, OG1RF_11113, or OG1RF_11123. However, structural modeling with Phyre2 [48] revealed that a portion of OG1RF_11123 has predicted structural homology to the channel-forming domain of colicin 1a [49] (Fig. S4D).

Orphan toxins encoded by a secretion system in a given strain can often be found as full-length toxins in other bacteria [45, 46]. Therefore, we used the sequences of OG1RF_11111, OG1RF_11113, and OG1RF_11123 as input for NCBI Protein BLAST to determine whether the orphan toxins we identified in *E. faecalis* OG1RF were found in other T7SS loci. We identified homologs to these orphan toxins in other *E. faecalis* strains as well as *Listeria sp*. (Fig. S4A-C, gray shading indicates regions of homology). These homologs were longer than the *E. faecalis* OG1RF genes and encoded N-terminal LXG domains, suggesting that in *Listeria* and other *E. faecalis* strains, homologs to OG1RF_11111, 11113, and 11123 are full-length toxins that could be secreted by the T7SS.

Interestingly, we identified an additional LXG gene product, OG1RF_12414, in a distal locus that is again notably absent from *E. faecalis* V583 (Fig. S5A and S5B). OG1RF_12414 has predicted structural homology to Tne2, a T6SS effector with NADase activity from *Pseudomonas protegens* (Fig. S5C) [50]. Additionally, we identified numerous C-terminal domains in LXG proteins distributed throughout the enterococci (Fig. S6). These include EndoU and Ntox44 nuclease domains [43, 51, 52], which have been characterized in effectors produced by other polymorphic toxin systems.

Polymorphic toxins are genetically linked to cognate immunity proteins that neutralize antagonistic activity and prevent self-intoxication [43, 52, 53]. Each of the five putative toxins in the OG1RF T7SS locus is encoded directly upstream of a small protein that could function in immunity. Whitney *et al*. demonstrated that the cytoplasmic antagonistic activity of *S. intermedius* LXG toxins TelA and TelB in *Escherichia coli* can be rescued by co-expression of cognate immunity factors [27]. Therefore, we examined if OG1RF_11110, 11112, 11122, or 12413 confer immunity to *E. faecalis* Δ*pip*_V583_ during phage infection of *E. faecalis* OG1RF. Constitutive expression of OG1RF_11122, and not OG1RF_11110, 11112, or 12413, partially neutralized phage induced T7SS antagonism (Fig. 2B), confirming an essential role for the OG1RF_11122 gene product in immunity, and suggesting that OG1RF_11121 is at least partly responsible for T7SS mediated intra-species antagonism. However, further investigation is needed to confirm whether the candidate immunity factors, OG1RF_11110, 11112 or 12413, are stably expressed under these experimental conditions.

To determine the contribution of OG1RF_11121 on intra- and interbacterial antagonism during phage infection, we measured the viability of phage resistant *E. faecalis* Δ*pip*_V583_ and *S. aureus* in co-culture with an *E. faecalis* OG1RF variant carrying a transposon insertion in OG1RF_11121 (OG1RF_11121-Tn). OG1RF_11121-Tn is equally susceptible to phage VPE25 infection compared to wild type *E. faecalis* OG1RF (Fig. S2A). Similar to the Δ*essB* (pCIEtm) strain carrying empty pCIEtm plasmid, phage infected OG1RF_11121-Tn (pCIEtm) did not inhibit the growth of the bystander bacteria (Fig. 2C – 2D). We were unable to clone OG1RF_11121 by itself into the inducible plasmid pCIEtm, suggesting leaky expression of OG1RF_11121 is toxic. Therefore, to complement *E. faecalis* OG1RF_11121-Tn we cloned both the OG1RF_11121 toxin and OG1RF_11122 antitoxin pair under the cCF10-inducible promoter in pCIEtm. Expression of both of these genes in the transposon mutant restored its ability antagonize T7SS susceptible bystanders (Fig. 2C – D). These data strongly suggest that the OG1RF_11121 encoded LXG toxin drives *E. faecalis* T7SS mediated antagonism of bystanders following phage infection.

### Sub-lethal antibiotic stress promotes T7SS dependent antagonism

Considering two genetically distinct phages trigger the induction of T7SS genes in *E. faecalis* [24], we reasoned that T7SS induction could be a result of phage mediated cellular damage and not specifically directed by a phage encoded protein. Antibiotics elicit a range of damage induced stress responses in bacteria [54–56]; therefore, independent of phage infection we investigated the effects of subinhibitory concentrations of antibiotics on T7SS expression in *E. faecalis*.

To investigate the influence of sublethal antibiotic concentrations on *E. faecalis* OG1RF T7SS transcription, we determined the minimum inhibitory concentrations (MIC) of ampicillin, vancomycin, and daptomycin (Fig.S7A – S7C) and monitored T7SS gene expression in *E. faecalis* OG1RF cells treated with a sub-lethal dose of antibiotic (50% of the MIC). We found that bacterial T7SS genes were significantly upregulated in the presence of the cell membrane targeting antibiotic, daptomycin, relative to the untreated control (Fig. 3A). In contrast, the cell wall biosynthesis inhibitors ampicillin and vancomycin either did not induce or had a minor impact on T7SS mRNA levels, respectively (Fig.3A). Additionally, induction of T7SS transcription occurred when bacteria were challenged with sub-inhibitory concentrations of the DNA targeting antibiotics ciprofloxacin and mitomycin C (Fig. 3B, Fig. S7D – S7E).

**Figure 3.**
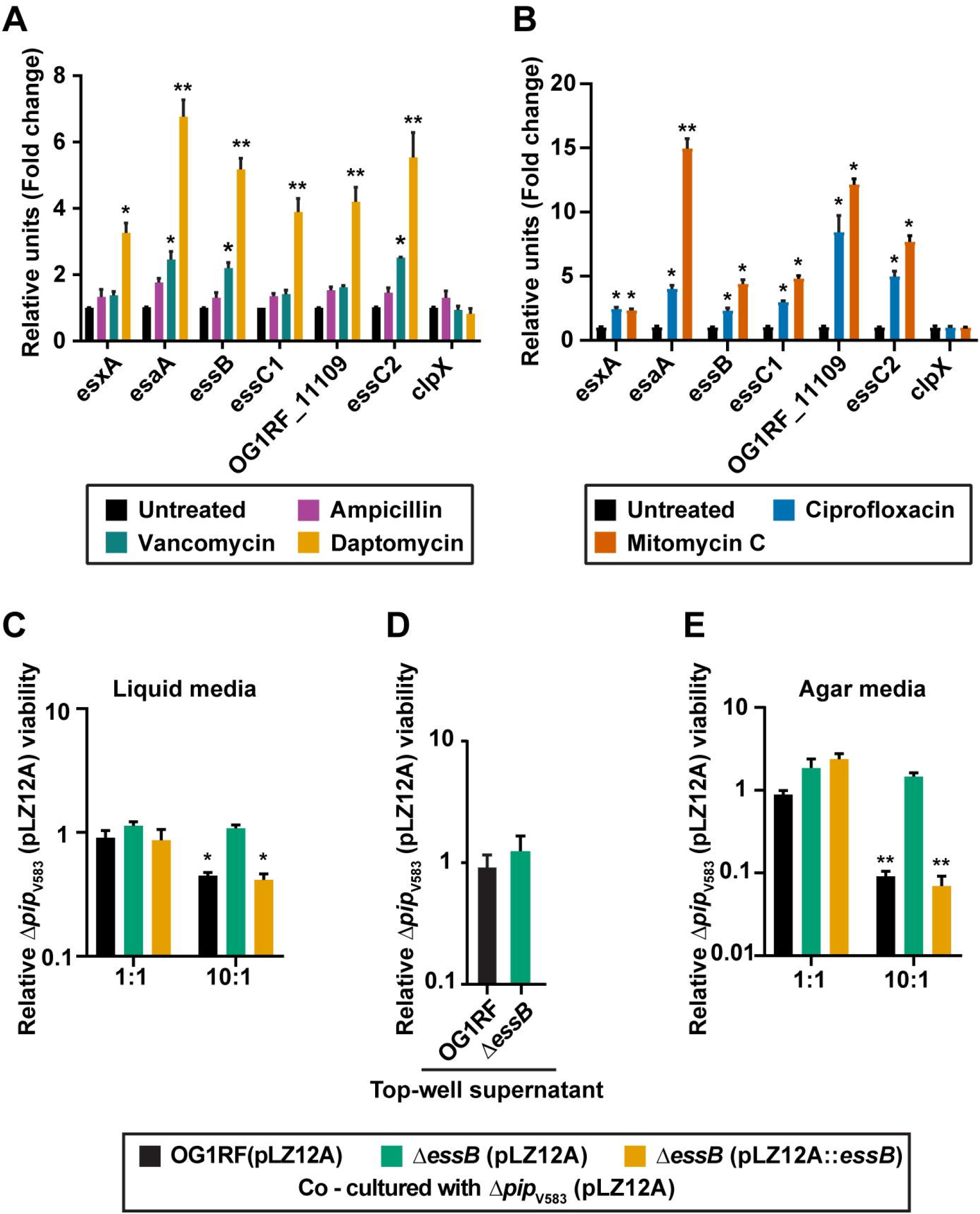
Sub-lethal antibiotic treatment enhances T7SS gene expression leading to inhibition of bystander bacteria. Altered expression of T7SS genes upon exposure to sub-inhibitory concentrations of **(A)** ampicillin (0.19 μg/ml), vancomycin (0.78 μg/ml) or daptomycin (6.25 μg/ml) and **(B)** ciprofloxacin (2 μg/ml) or mitomycin C (4 μg/ml) for 40 minutes relative to the untreated control. *clpX* is shown as a negative control. **(C-E)** Contact–dependent T7SS mediated inhibition of bystander bacteria in the presence of daptomycin. Relative viability of *E. faecalis* Δ*pip*_V583_ was measured during co-culture with *E. faecalis* OG1RF or Δ*essB* antagonists in the presence and absence of daptomycin treatment in **(C)** liquid culture (2.5 μg/ml daptomycin), **(D)** trans-well plates to prevent physical engagement between cells (2.5 μg/ml daptomycin) and **(E)** in contact on agar media (0.5 μg/ml daptomycin). Δ*essB* (pLZ12A) and Δ*essB* (pLZ12A::*essB*) represent the empty vector control and complemented strains. Data show three biological replicates. Error bars indicate standard deviation. \**P* < 0.01, \*\**P* < 0.001 to 0.0001 by unpaired Student’s t-test.

We next sought to assess the influence of daptomycin driven T7SS induction on inter-enterococcal antagonism. *E. faecalis* V583 and its derivatives are more sensitive to daptomycin compared to *E. faecalis* OG1RF strains (Fig. S8A – S8C), so we applied a reduced concentration of 2.5 μg/ml daptomycin in the co-culture inhibition assay to prevent daptomycin intoxication of *E. faecalis* Δ*pip*_V583_ bystanders. Because *E. faecalis* OG1RF T7SS gene expression is less robust in the presence of 2.5 μg/ml compared to 6.25 μg/ml daptomycin, which was used in our previous experiments (Fig. 3A and S8D), a 10:1 ratio of daptomycin treated *E. faecalis* OG1RF was required for growth inhibition of *E. faecalis* Δ*pip*_V583_ during co-culture (Fig. 3C). Consistent with our previous results, daptomycin induced T7SS inhibition of *E. faecalis* Δ*pip*_V583_ was contact dependent (Fig. 3D). To increase T7SS mediated contact-dependent killing of the target strain during daptomycin exposure, we performed the inhibition assay on nutrient agar plates. The sub-inhibitory concentration of daptomycin (2.5 μg/ml) used in liquid culture was toxic to the cells on agar plates (Fig. S8E), so we lowered the daptomycin concentration to 0.5 μg/ml to prevent drug toxicity in the agar-based antagonism assay. Plating T7SS producing *E. faecalis* OG1RF cells and *E. faecalis* Δ*pip*_V583_ bystander cells at a ratio of 10:1 resulted in ~10–fold inhibition of bystander growth (Fig. 3E). Although 0.5 μg/ml of daptomycin did not dramatically increase *E. faecalis* OG1RF T7SS transcript abundances, this was sufficient to promote daptomycin mediated T7SS inhibition of bystanders on agar plates (Fig. 3E and Fig. S8E). These data show that in addition to phages, antibiotics can be sensed by *E. faecalis* thereby inducing T7SS antagonism of non-kin bacterial cells. These data also show that the magnitude of T7SS gene expression and forcing bacteria-bacteria contact is directly related to the potency of T7SS inhibition.

### The primary bile acid sodium cholate does not modulate *E. faecalis* T7SS gene expression

To gain insight into host-associated environmental cues that could trigger *E. faecalis* OG1RF T7SS, we measured T7SS transcription in the presence of a sub-inhibitory concentration of the primary bile acid sodium cholate, an abundant compound found in the mammalian intestinal tract and that is known to promote bacterial cell membrane stress [57, 58]. 4% sodium cholate, a concentration that has been shown to severely impair the growth of *E. faecalis* OG1RF cell envelop mutants, caused only a minor reduction in cell density of wild type *E. faecalis* OG1RF [59] (Fig. S9A) and it did not stimulate T7SS gene expression (Fig. S9B). Collectively, these data show that T7SS induction in *E. faecalis* occurs in response to select cell envelope stressors.

### IreK and OG1RF_11099 facilitate T7SS expression in phage infected *E. faecalis* OG1RF via a non-canonical signaling pathway

Having established that both phage and daptomycin mediated membrane damage independently stimulates heightened *E. faecalis* OG1RF T7SS gene expression and antagonistic activity, we next sought to identify the genetic determinants that sense this damage and promote T7SS transcription. Two-component systems, LiaR/S and CroS/R, and the PASTA kinase family protein IreK are well-characterized modulators of enterococcal cell envelope homeostasis and antimicrobial tolerance [60–62]. Aberrant cardiolipin microdomain remodeling in the bacterial cell membrane in the absence of the LiaR response regulator results in daptomycin hypersensitivity and virulence attenuation [63]. CroS/R signaling and subsequent modulation of gene expression govern cell wall integrity and promote resistance to cephalosporins, glycopeptides and beta—lactam antibiotics [64–66]. The *ireK* encoded transmembrane Ser/Thr kinase regulates cell wall homeostasis, antimicrobial resistance, and contributes to bacterial fitness during long-term colonization of the intestinal tract [61, 67, 68]. Recently it has been shown that direct cross-talk between IreK and the CroS/R system positively impacts enterococcal cephalosporin resistance [69].

Wild type *E. faecalis* OG1RF, an *ireK* in-frame deletion mutant [61] and transposon (Tn) insertion mutants of *liaR*, *liaS*, *croR*, and *croS* [70] all display similar growth kinetics in the absence of phage VPE25 infection (Fig. S10A). Although *croR*-Tn and *croS*-Tn exhibit reductions in the plaquing efficiency of VPE25 particles, none of these genetic elements of enterococcal cell wall homeostasis and antibiotic resistance were required for VPE25 infection (Fig. S10B). We queried the expression levels of T7SS genes in these isogenic mutants during phage VPE25 infection (MOI = 1). T7SS gene expression was not enhanced in the Δ*ireK* mutant during phage infection (Fig. 4A), whereas *liaR*-Tn, *liaS*-Tn, *croR*-Tn, and *croS*-Tn produced heightened levels of T7SS transcripts similar to the wild type *E. faecalis* OG1RF compared to the uninfected controls (Fig. S11A – S11F). A sub-lethal concentration of the cephalosporin ceftriaxone did not induce T7SS gene expression (Fig. S12A), indicating that expression of T7SS genes following phage mediated membrane damage signals through a pathway that is distinct from the IreK response to cephalosporin stress. Additionally, the Δ*ireK* mutant phenocopies the Δ*essB* mutant strain in the interbacterial antagonism co-culture assay, wherein the Δ*ireK* mutant is unable to mediate phage induced T7SS dependent killing of the phage resistant *E. faecalis* Δ*pip*_V583_ non-kin cells (Fig. 4B). T7SS antagonism is restored in *E. faecalis* Δ*ireK* by introducing the wild type gene in *trans* (Fig. 4B). Collectively, these results indicate that IreK senses phage mediated membrane damage promoting T7SS transcription independent of the CroS/R pathway.

**Figure 4.**
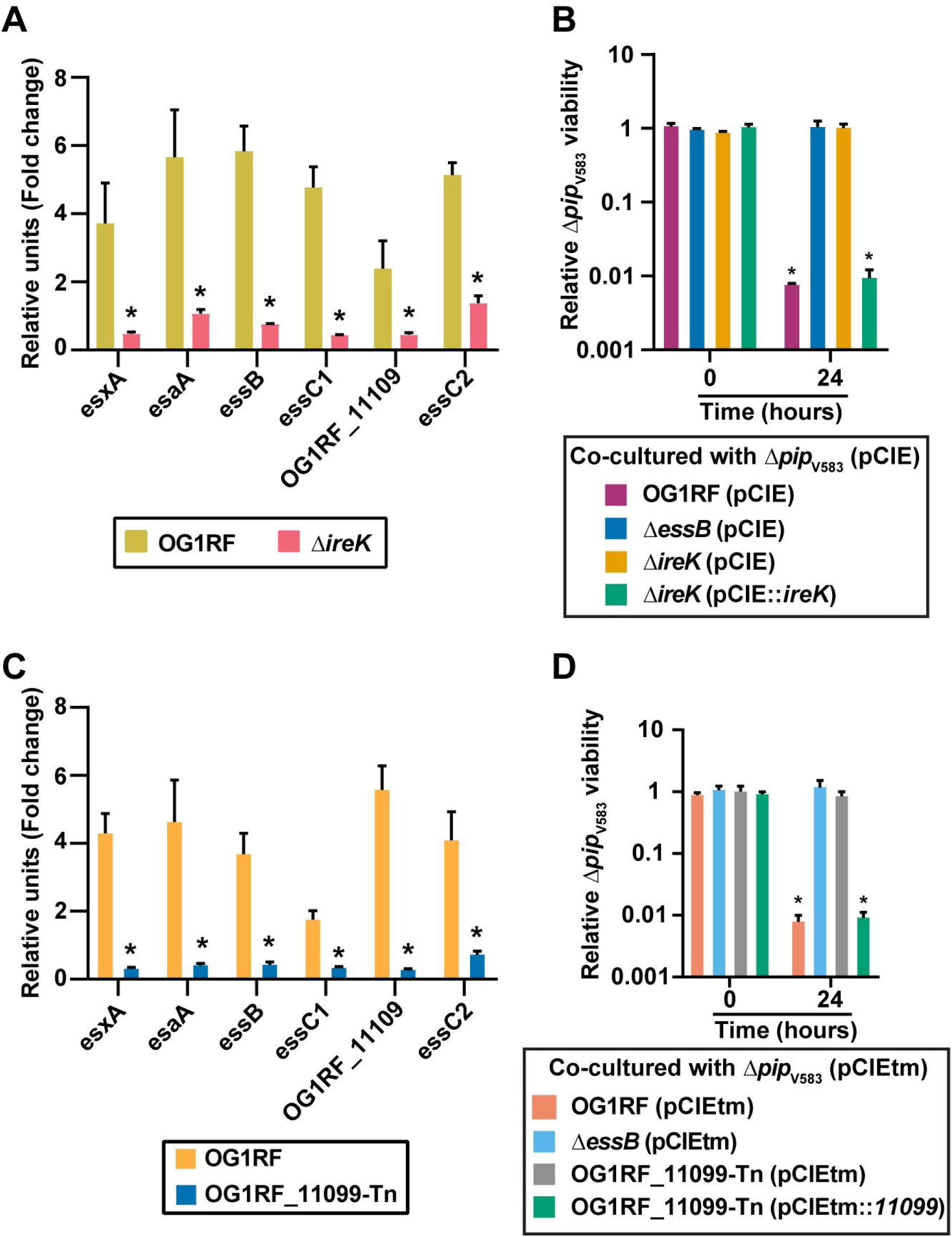
IreK and OG1RF_11099 control transcription of enterococcal T7SS genes and subsequent inhibition of bystander bacteria during phage infection. **(A)** Phage infection leads to enhanced expression of T7SS genes in wild type *E. faecalis* OG1RF but not in a Δ*ireK* mutant strain. **(B)** Growth inhibition of *E. faecalis* Δ*pip*_V583_ during phage infection of *E. faecalis* OG1RF is abrogated in the Δ*essB* and Δ*ireK* mutants carrying empty pCIEtm. pCIEtm::*ireK* complemented the T7SS activity defect of the Δ*ireK* strain. **(C)** Disruption of OG1RF_11099 leads to reduced expression of T7SS genes during phage infection. The data are represented as the fold change of normalized mRNA relative to uninfected samples at the same time points. **(D)** T7SS dependent intraspecies antagonism during phage infection is alleviated in the presence of OG1RF_11099-Tn mutant carrying empty pCIEtm. pCIEtm::*11099* complemented the T7SS activity defect of the OG1RF_11099-Tn mutant strain. Data represent three biological replicates. Error bars indicate standard deviation. \**P* < 0.00001 by unpaired Student’s t-test.

OG1RF_11099, located immediately upstream of the T7SS cluster is predicted to encode a GntR family transcriptional regulator, thus we sought to assess the contribution of OG1RF_11099 on T7SS transcription and functionality. *E. faecalis* OG1RF carrying a transposon insertion in OG1RF_11099 is equally susceptible to phage VPE25 infection compared to wild type *E. faecalis* OG1RF (Fig. S2A) In contrast to wild type *E. faecalis* OG1RF, T7SS genes were not induced during phage predation of *E. faecalis* OG1RF_11099-Tn (Fig. 4C). We evaluated the influence of OG1RF_11099-dependent regulation on the activity of T7SS in intraspecies antagonism using our co-culture assay. Similar to *E. faecalis* Δ*essB*, the OG1RF_11099-Tn mutant displayed attenuated T7SS activity in phage infected co-cultures (Fig. 4D). T7SS dependent antagonism of *E. faecalis* OG1RF_11099-Tn could be restored following complementation (Fig. 4D). Collectively, these results indicate that *OG1RF_11099* encodes a positive regulator of *E. faecalis* T7SS important for phage mediated inhibition of bystander bacteria. Given that IreK governs downstream signaling events via phosphorylation [71], and the fact that OG1RF_11099 was not differentially expressed in response to phage infection of wild type *E. faecalis* OG1RF or *ireK* mutant strains (Fig. S12B), suggests that either post-translational modification of OG1RF_11099 or a yet unidentified protein downstream of IreK engaging with OG1RF_11099 accounts for T7SS gene expression during phage infection.

## Discussion

Despite the fact that bacteria exist in complex microbial communities that socially interact [72, 73], phage predation studies have primarily been performed in monoculture [24, 74–76]. Studies report phage-mediated effects on non-target bacteria linked to interbacterial interactions and evolved phage tropism for non-cognate bacteria [77–79], whereas other studies have identified minimal changes in microbiota diversity during phage therapy [77, 80].

Our results extend previous work that observed the induction of *E. faecalis* OG1RF T7SS gene expression in response to phage infection [24]. By using an *in vitro* antibacterial antagonism assay, we discovered that phage predation of *E. faecalis* OG1RF has an inhibitory effect on non-phage targeted bacterial species during co-culture. Our work shows that phage mediated inhibition of Gram positive bystander bacteria relies on the expression and activity of T7SS genes. This work establishes a framework to begin investigating if and how phage infection of target bacteria influences non-target bacterial populations in complex communities such as the microbiota.

Our data suggest that membrane stress associated with phage infection or sub-lethal daptomycin treatment stimulates T7SS mediated antibacterial antagonism of *E. faecalis* OG1RF (Fig. 5). Given that daptomycin is used to target vancomycin-resistant enterococcal infections, this finding provides a hypothesis for how antibiotic-resistant enterococci achieve overgrowth and dominate the microbiota following antibiotic treatment. Further investigation is required to understand how T7SS induction might contribute to enterococcal fitness in polymicrobial environments. Although exposure to a sub-inhibitory level of primary bile salt (a common molecule found in the intestine) did not elicit T7SS expression, it is possible that other stressors encountered in the intestinal tract, including lysozyme, antimicrobial proteins, and nutrient availability could influence T7SS activity in *E. faecalis*. Indeed, *E. faecalis* T7SS mutants are defective in their ability to colonize the murine reproductive tract, which like the intestine is a polymicrobial environment [36].

**Figure 5.**
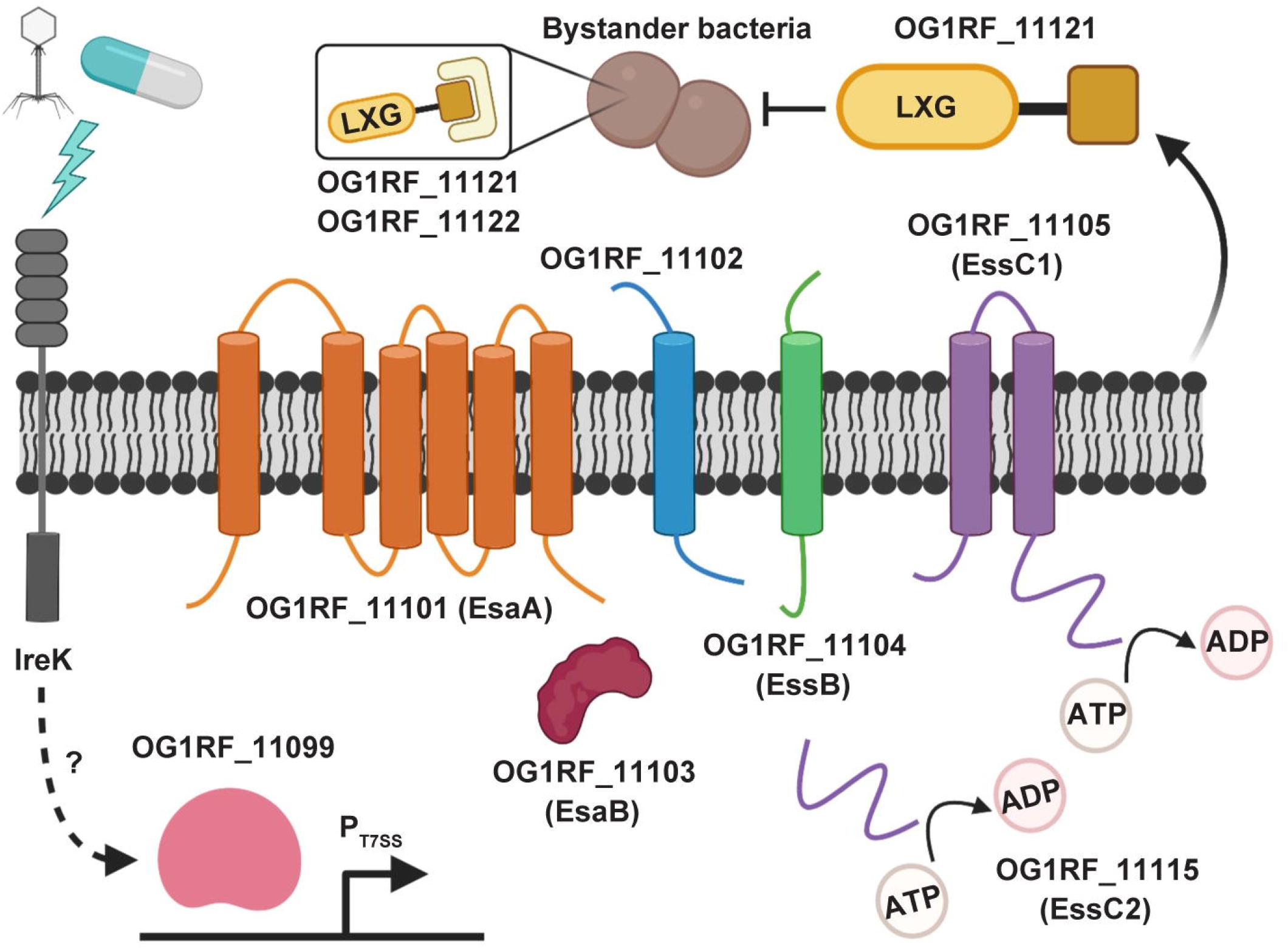
A model for inhibition of bystander bacteria by the *E. faecalis* OG1RF T7SS. Phage and select antibiotics trigger a response involving IreK that results in induction of expression of T7SS genes. Transcription of T7SS genes is regulated by the predicted GntR-family transcription factor OG1RF_11099. The predicted core components of the OG1RF T7SS machinery are putative membrane proteins EsaA (OG1RF_11101), OG1RF_11102, EssB (OG1RF_11104), and EssC1 (OG1RF_11105) as well as the putative cytoplasmic protein EsaB (OG1RF_11103). EssC2 (OG1RF_11115) lacks transmembrane domains and is thus not predicted to be membrane-anchored. Upon induction of the T7SS, OG1RF_11121 is secreted from the cell, resulting in antibacterial activity against select neighboring bacteria. Expression of OG1RF_11122 can partially block toxicity caused by OG1RF_11121. Predictions of membrane topology were obtained using TMHMM [108]. The figure was created with Biorender.com.

We discovered that transcriptional activation of the T7SS during phage infection relies on IreK (Fig. 5). Previously characterized IreK–mediated stress response pathways, including cephalosporin stress or CroS/R signaling, did not contribute to T7SS expression. We hypothesize that IreK senses diverse environmental stressors and coordinates distinct outputs in response to specific stimuli. Considering that IreK signaling is important for *E. faecalis* intestinal colonization [68], it is possible that IreK–dependent T7SS expression in response to intestinal cues modulate interbacterial interactions and enterococcal persistence in the intestine. However, the molecular mechanism by which IreK facilitates T7SS transcription remains unanswered. Additionally, we currently do not know if IreK directly senses phage or daptomycin mediated membrane damage or some other signal feeds into IreK to facilitate T7SS induction.

Additionally, we discovered that *E. faecalis* OG1RF T7SS transcription is regulated by a GntR-family transcriptional regulator encoded by OG1RF_11099, a gene found immediately upstream of the T7SS cluster (Fig. 5). Interestingly, OG1RF_11099 is highly conserved across enterococci, including *E. faecalis* V583 (Fig. 1A) and other strains that lack T7SS. The presence of a conserved transcriptional regulator in the absence of its target genetic region supports the idea that certain strains of enterococci have undergone genome reduction as an evolutionary strategy to adapt to unique host and non-host environments. It is possible that in *E. faecalis* V583, the OG1RF_11099 homolog (EF1328) has been retained to regulate other genes within the regulon that are less dispensable than T7SS. Additionally, our data indicate that OG1RF_11099 transcription is not dependent on IreK or and is not induced during phage infection of wild type *E. faecalis* OG1RF. Previously published work demonstrated that IreK kinase activity is essential for driving the cell wall stress response in *E. faecalis* [67, 71]. Therefore, we hypothesize that IreK directly or indirectly regulates OG1RF_11099 activity for T7SS expression via post-translational modification.

Antibacterial properties of T7SS substrates have been demonstrated [26, 27, 40]. Here we provide evidence that mutation in the LXG toxin encoded by OG1RF_11121 abrogates phage induced T7SS dependent inhibition of bystander bacteria while expression of the downstream immunity gene OG1RF_11122 in T7SS targeted *E. faecalis* Δ*pip*_V583_ cells conferred partial protection from this inhibition. It is possible that constitutive expression of OG1RF_11122 from a multicopy plasmid results in elevated accumulation of OG1RF_11122 in the bystander strain which is toxic and could account for the partial protection phenotype. Aside from its LXG domain, OG1RF_11121 does not harbor any other recognizable protein domains, hence the mechanism underlying its toxicity is unclear. Whitney *et al*. demonstrated that LXG toxin antagonism is contact–dependent, having minimal to no impact on target cells in liquid media [27]. Although we found that physical engagement is crucial for *E. faecalis* T7SS mediated antagonism, we observed a significant reduction in target cell growth in liquid media both during phage and daptomycin treatment of T7SS proficient *E. faecalis*.

In contrast to the broad antagonism of *S. intermedius* T7SS [27], the *E. faecalis* OG1RF T7SS targets a more limited number of bacterial species. Interestingly, *E. faecalis* OG1RF T7SS antagonism is ineffective against various species of streptococci, which like the enterococci are lactic acid bacteria. Nucleotide– and protein–based homology searches did not reveal homologs of candidate immunity proteins, OG1RF_11110, OG1RF_11112, OG1RF_11122, or OG1RF_12413, in *S. agalactiae* COH1. Genome sequences of the other four streptococci used in this present study are not available, and hence we cannot comment on the presence of potential immunity proteins against OG1RF T7SS toxins in these strains. However, resistance of multiple streptococcal species to OG1RF T7SS mediated inhibition suggest that common cell surface modifications, e.g., capsule or surface polysaccharides, might be responsible for blocking toxin activity. Narrow target range is a common attribute of contact-dependent toxins that interact with specific membrane receptors on target cells to exert inhibitory activity [81]. However, specific receptors of T7SS toxins are yet to be identified. It is possible that specific or non-specific interactions between the *E. faecalis* OG1RF and *S. aureus* or *L. monocytogenes* cell surfaces facilitate T7SS interbacterial antagonism and such interactions are incompatible or occluded for the streptococci.

It is currently unknown whether T7SS toxin delivery requires contact with a receptor on target cells or whether delivery can occur in the absence of a receptor. Examples of both methods of toxin delivery are widespread in bacteria. Toxins such as colicins and R-pyocins mediate contact with target cells via protein receptors and LPS, respectively [82–84]. Delivery of colicins and toxins produced by contact-dependent inhibition systems in Gram-negative bacteria requires interactions with receptors at the outer and inner membranes [85, 86]. Conversely, the T6SS needle-like machinery that punctures target cell envelopes delivers toxins in a contact-dependent, receptor-independent manner [87]. Cell surface moieties can also affect recognition of target cells and subsequent toxin delivery. The presence of capsule can block target cell recognition by contact-dependent growth inhibition systems in *Acinetobacter baumannii* [88], *E. coli* [89] and *Klebsiella pneumoniae* [90]. Therefore, it is possible that a feature of the streptococcal cell surface, such as capsule modifications, renders them insensitive to killing by toxins delivered by the *E. faecalis* OG1RF T7SS.

Enterococci occupy polymicrobial infections often interacting with other bacteria [91–94]. Although commensal *E. faecalis* antagonize virulent *S. aureus* through the production of superoxide [95], the two species also exhibit growth synergy via exchange of critical nutrients [96]. Here, we show that phage treatment of *E. faecalis* OG1RF can indirectly impact the growth of neighboring phage-resistant bacteria, including *S. aureus*, in a T7SS–dependent manner, suggesting that phage therapy directed against enterococci and driving T7SS activity could be useful for the treatment of polymicrobial infections. However, the counter argument is that phage therapy directed against enterococci could push a bacterial community toward dysbiosis, as phage induced T7SS activity could directly inhibit beneficial bystander bacteria. This raises questions about the consequences of phage mediated off-target effects on bacteria. Could phage induced T7SS activity be used to reduce phage expansion into other closely related strains as a means to dilute phages out of a population, or is it simply that phage induction of the T7SS serves as a mechanism that benefits a select few within a population to aid in their reoccupation of a niche upon overcoming phage infection? Future studies aimed at exploring enterococcal T7SS antagonism in polymicrobial communities should help elucidate the impact of phages on microbial community composition.

## Materials and Methods

### Bacteria and bacteriophages

Bacteria and phages used in this study are listed in Table S1. Bacteria were grown with aeration in Todd-Hewitt broth (THB) or on THB agar supplemented with 10mM MgSO_4_ at 37°C. The following antibiotic concentrations were added to media for the selection of specific bacterial strains or species: *E. faecalis* OG1RF (25◻μg/ml fusidic acid, 50 μg/ml rifampin), *E. faecalis* V583 Δ*pip*_V583_ (25 μg/ml or 100◻μg/ml gentamicin in liquid and agar media, respectively), *S. aureus* AH2146 LAC Φ11:LL29 (1◻μg/ml tetracycline), *L. monocytogenes* 10403S (100◻μg/ml streptomycin), *S. gordonii* ATCC 49818 (500 μg/ml streptomycin), *S. salivarius* K12 (100 μg/ml spectinomycin), *V. cholerae* C6706 int I4::TnFL63 and *S. enterica* serovar Typhimurium 140285 put::Kan (50◻μg/ml kanamycin). *S. agalactiae* COH1 was distinguished from *E. faecalis* on Chrome indicator Agar (CHROMagar StrepB SB282). We were unable to differentially select *E. coli, L. lactis*, *S. pyogenes* and *S. mitis* from *E. faecalis* based on antibiotic sensitivity. Therefore, colony counts of these bacteria in co-culture experiments were acquired by subtracting the *E. faecalis* colony numbers on selective media from the total number of colonies on non-selective media. Strains harboring pLZ12A and its derivatives were grown in the presence of 20◻μg/ml chloramphenicol and strains carrying pCIEtm and pCIEtm derivatives were selected on media containing 5 μg/ml tetracycline.

### Bioinformatic analyses

Genome sequences of *E. faecalis* V583 (NC_004668.1) and OG1RF (NC_017316.1) were obtained from NCBI. Alignments were generated and visualized using EasyFig [97]. OG1RF protein domains were identified using KEGG [98] and ExPASy PROSITE [99]. Structure modeling of OG1RF_12414 was done with Phyre2 [48]. Crystal structures overlays were generated using Pymol [100]. The EsxA phylogenetic tree was constructed in MEGA version X [101] using non-redundant protein sequences obtained from NCBI BLAST [102] with OG1RF_11100 as input and was edited using the Interactive Tree Of Life browser [103]. OG1RF_11109 was used as an input for the NCBI Conserved Domain Architecture Retrieval Tool [104] to identify protein domains that co-occur with LXG domains in *Enterococcus* (NCBI:txid1350).

### Antibiotic sensitivity profiles

Antibiotic susceptibility profiles for ampicillin, vancomycin, and daptomycin were determined using a broth microdilution assay. Overnight (O/N) *E. faecalis* OG1RF cultures were diluted to 5◻×◻10^6^ CFU/ml and 100 μl was added to each well of a 96-well plate to give a final cell density of 5◻×◻10^5^ CFU/ml. Antibiotic stocks were added to the first column of each row, mixed thoroughly, and serially diluted 2-fold across the rows. The last column was used as a no drug control. Cultures containing daptomycin were supplemented with 50◻μg/ml CaCl_2_. Bacterial growth was monitored by measuring absorbance (OD_600_) using a Synergy H1 microplate reader set to 37°C with continuous shaking O/N. Growth curves are presented as the average of three biological replicates. A concentration of antibiotic just below the drug amount that inhibits bacterial growth was deemed sub-lethal and used to examine T7SS genes expression.

### Co-culture bacterial antagonism assays

For inter- and intraspecies antagonism assays in liquid media, O/N cultures of different bacteria were diluted in THB containing 10mM MgSO_4_ to an OD_600_ of 0.2 and mixed together in a 1:1 or 10:1 ratio. The mixed cell suspensions were either left untreated or treated with phage VPE25 (MOI 0.01) or daptomycin (2.5 μg/ml) and grown at 37°C with aeration. For pheromone induction of genes OG1RF_11121, *ireK* and OG1RF_11099 cloned into pCIEtm, 10 ng/ml cCF10 (from Mimotopes) was added at the time of phage administration. For antagonism experiments on agar plates, O/N cultures of different strains were diluted to an OD_600_ of 0.2 and mixed together in a 1:1 or 10:1 ratio. A total of 10^7^ cells from mixed culture suspension was added to 5 ml THB + 0.35% agar at 55°C and were poured over the surface of a THB agar plate in the absence or presence of daptomycin (0.5 μg/ml). The plates were incubated at 37°C under static conditions for 24 hours. Cells were harvested by scraping off the top agar, resuspending in 5 ml of PBS, and the cfus were obtained by plating serially diluted cell suspension on appropriate selective agar plates. Relative viability was calculated from the ratio of target strain cfu in the treated versus the untreated co-culture. The assays were performed in biological triplicates.

### RNA extraction and quantitative PCR

RNA was extracted from phage, antibiotic, or 4% sodium cholate treated or untreated *E. faecalis* OG1RF cells using an RNeasy Mini Kit (Qiagen) with the following published modifications [24]. cDNA was generated from 1 μg of RNA using qScript cDNA SuperMix (QuantaBio) and transcript levels were analyzed by qPCR using PowerUp SYBR Green Master Mix (Applied Biosystems). Transcript abundances were normalized to 16S rRNA gene transcripts and fold–change was calculated by comparing to untreated controls. All data are represented as the average of three biological replicates. All the primers used for qPCR are listed in Table S1.

### Bacterial growth curves

25 ml of 10mM MgSO_4_ supplemented THB was inoculated with O/N cultures of *E. faecalis* diluted to an OD_600_ of 0.025 and distributed to a 96-well plate in 0.1 ml volumes. Cultures were incubated at 37° C with aeration. OD_600_ was measured periodically for 18 hours in a Synergy H1 microplate reader.

### Efficiency of plating (EOP) assays

To investigate if phage VPE25 can infect and lyse *E. faecalis* mutants and various other bacterial species, 10^7^ PFU/ml of phage was serially diluted and the phage was titered on each strain using a THB agar overlay plaque assay. EOP is expressed as the percentage of phage titer from each strain relative to the wild type *E. faecalis* OG1RF control. Data are presented as the average of three biological replicates.

### Construction of *E. faecalis* mutants and complementation

Isolation of *E. faecalis* genomic DNA was performed using a ZymoBIOMICS DNA Miniprep Kit (Zymo Research). All PCR used for cloning were performed with high fidelity KOD Hot Start DNA Polymerase (EMD Millipore). *E. faecalis* Δ*essB* was generated by allelic replacement by cloning an in frame *essB* deletion product into pLT06 using Gibson Assembly^®^ Master Mix (New England Biolabs), integrating this construct into the chromosome, and resolving the deletion mutant by homologous recombination [105–107]. For ectopic expression of putative immunity proteins, coding regions of OG1RF_11110, OG1RF_11112, OG1RF_11122, and OG1RF_12413 were cloned downstream of the *bacA* promoter (P_bacA_) by restriction digestion and ligation into the shuttle vector pLZ12A [20]. Coding regions of *ireK* and OG1RF_11099 were cloned downstream of the cCF10 responsive promoter (P_Q_) by restriction digestion and ligation into pCIE and pCIEtm vectors, respectively. As attempts to clone OG1RF_11121 by itself were unsuccessful, we cloned the OG1RF_11121 and OG1RF_11122 open reading frames, which overlap by 13 base pairs, together under the P_Q_ promoter in pCIEtm plasmid. Primer sequences and restriction enzymes used for cloning are listed in Table S1. Plasmids were introduced into electrocompetent *E. faecalis* cells as previously described [20].

### Statistical analysis

Statistical tests were performed using GraphPad – Prism version 8.2.1. For qPCR and bacterial competition assays, unpaired Student’s t-tests were used. *P* values are indicated in the figure legends.

### Data availability

All raw data are available upon request.

## Acknowledgments

This work was supported by National Institutes of Health grants R01AI141479 (B.A.D.) and R01AI122742 (G.M.D.). J.L.E.W. was supported by American Heart Association Grant 19POST34450124 / Julia Willett / 2018. We would like to thank Andrés Vázquez-Torres, Laurel Lenz, Alex Horswill, Stefan Pukatzki, Kelly Doran, and their lab members for sharing bacterial strains used in this study. We thank Michelle Korir for the *ireK* complementation plasmid construct and strain.

## Author Contributions

A.C., J.L.E.W., G.M.D. and B.A.D. designed the study. A.C. and J.L.E.W. performed experiments and bioinformatic analyses. A.C., J.L.E.W., G.M.D. and B.A.D. analyzed data. A.C., J.L.E.W. and B.A.D. wrote the paper with input from G.M.D.

---

**Figure S1. Phylogenetic tree of EsxA sequences in *enterococci***. Non-redundant sequences (n=96) were identified using NCBI BLAST with OG1RF EsxA (OG1RF_11100) as the input. The tree was constructed in MEGAX using the Maximum Likelihood method and JTT matrix-based model and is drawn to scale, with branch lengths measured in the number of substitutions per site. The tree with the highest log likelihood (−3544.39) is shown. *E. faecalis* sequences are highlighted in purple, and the GenBank identifier for EsxA from OG1RF (AEA93787.1) is shown in red font.

**Figure S2. Phage VPE25 infects wild type *E. faecalis* OG1RF and T7SS mutants with similar efficacy**. **(A)** The measurement of phage particles released from wild type *E. faecalis* OG1RF, Δ*essB*, OG1RF_11121-Tn, and OG1RF_11099-Tn mutant strains following phage VPE25 infection. **(B)** Viability of strains of the OG1RF background exhibiting differential T7SS activity in the absence and presence of phage. **(C)** Viability of T7SS susceptible strains during intraspecies competition experiments in the absence and presence of phage. Data represent three biological replicates. Error bars indicate standard deviation. \**P* < 0.0001 by unpaired Student’s t-test.

**Figure S3. Phage induced T7SS inhibitory activity is contact dependent**. Intraspecies competition experiment performed in the presence of unfiltered supernatant from phage treated and untreated *E. faecalis* wild type OG1RF or Δ*essB* added **(A)** to the top of a well separated by a 0.4 μm membrane from the bottom well containing *E. faecalis* Δ*pip*_V583_ culture, and bacterial viability was determined after 24 hours, or **(B)** directly into *E. faecalis* Δ*pip*_V583_ culture in microtiter plate wells (*P* = 0.7955 by two-way analysis of variance [ANOVA]). **(C)** Growth of Δ*pip*_V583_ was monitored in the presence of filtered supernatant from uninfected and phage infected cultures of wild type *E. faecalis* OG1RF and Δ*essB* (*P* = 0.0883 by two-way analysis of variance [ANOVA]). *E. faecalis* Δ*pip*_V583_ cultures in all of these three contact-dependent assays contained gentamicin (25◻μg/ml) to prevent growth of the OG1RF background strains that may have carried over in unfiltered supernatants. Error bars indicate standard deviation.

**Figure S4. Putative orphan toxins in OG1RF are found as full-length LXG-domain proteins in other bacteria**. OG1RF_11111, OG1RF_11113, and OG1RF_11123 sequences were used as input for NCBI BLAST. Alignments and homology were rendered in EasyFig.

**Figure S5. A distal *E. faecalis* OG1RF locus encodes an additional LXG-domain protein**. **(A)** Schematic showing homology between V583 (NC_004668.1, top) and OG1RF (NC_017316.1, bottom). Sequences were obtained from NCBI, and homology comparisons were rendered in EasyFig. **(B)** Cartoon depicting the LXG domain of OG1RF_12414 (identified using KEGG and ExPASy PROSITE). **(C)** Predicted structural homology between OG1RF_12414 (lilac) and the *Pseudomonas protogens* Pf-5 Tne2/Tni2 complex (PDB 6B12). Tne2 is shown in green, and Tni2 is shown in gray. Structural modeling was done using PHYRE2, and images were rendered in Pymol.

**Figure S6. Domain architecture of enterococcal LXG proteins**. Domain architectures were identified using the NCBI Conserved Domain Architectural Retrieval Tool (DART) with OG1RF_11109 as an input. Diagrams are drawn to scale.

**Figure S7. Antibiotic susceptibility of *E. faecalis* OG1RF**. Growth of wild type *E. faecalis* OG1RF was monitored over 20 hours in the presence or absence of **(A)** ampicillin, **(B)** vancomycin and **(C)** daptomycin in microtiter plates. The antibiotic concentrations highlighted with a blue box were deemed sub-inhibitory and used to investigate T7SS gene expression levels. Early log-phase cultures of *E. faecalis* OG1RF were grown in the presence or absence of **(D)** mitomycin C (4 μg/ml) or **(E)** ciprofloxacin (2 μg/ml) to show that these concentrations of DNA targeting antibiotics do not prevent bacterial growth. Error bars indicate standard deviation.

**Figure S8. Impact of daptomycin concentration on *E. faecalis* growth and T7SS induction**. Growth of different enterococcal strains either untreated or treated with 6.25 μg/ml, 2.5 μg/ml or 0.5 μg/ml of daptomycin in **(A – C)** liquid media. **(D)** T7SS transcripts were measured from *E. faecalis* OG1RF cells grown in liquid media containing either no daptomycin or 6.25 μg/ml, 2.5 μg/ml, or 0.5 μg/ml of daptomycin. The data are expressed as the average of three biological replicates ± the standard deviation. *P* < 0.001 by unpaired Student’s t-test. **(E)** Viable bacterial cells recovered from growth on daptomycin supplemented agar media for 24 hours. The dashed line indicates the limit of detection.

**Figure S9. Effect of sub-lethal bile salt treatment on growth and T7SS transcription in *E. faecalis* OG1RF**. **(A)** Optical density of wild type *E. faecalis* OG1RF grown in the absence and presence of 4% sodium cholate was measured for 18 hours. **(B)** Transcript levels of OG1RF T7SS genes in untreated and 4% sodium cholate treated E. faecalis OG1RF after 4 hours. *P* < 0.001 to by unpaired Student’s t-test. Error bars indicate standard deviation.

**Figure S10. *E. faecalis* mutants of cell wall homeostasis show no growth defects and respond to phage VPE25 infection**. **(A)** Optical density of wild type *E. faecalis* OG1RF and isogenic mutants were monitored for 18 hours. **(B)** While all strains were susceptible to phage VPE25 infection, the proportion of released phage particles was diminished in the *croR* and *croS* transposon mutant background. Data represent three biological replicates. Error bars indicate standard deviation. \**P* < 0.001 by unpaired Student’s t-test.

**Figure S11. Quantitative PCR demonstrates that LiaR/S and CroS/R two-component systems do not influence T7SS gene expression during phage infection**. **(A-F)** mRNA transcript levels of T7SS genes are enhanced in the transposon mutants of *liaR*, *liaS*, *croR* and *croS* strains similar to wild type *E. faecalis* OG1RF during phage infection (MOI = 1) compared to untreated controls. Data represent three biological replicates. Error bars indicate standard deviation. \**P* < 0.01, \*\**P* < 0.0001 by unpaired Student’s t-test.

**Figure S12. Influence of sub-lethal ceftriaxone challenge and phage infection on the expression of *E. faecalis* T7SS genes**. **(A)** Transcription of T7SS genes in *E. faecalis* OG1RF are not elevated 20 minutes post ceftriaxone (128μg/ml) administration relative to an untreated control. **(B)** OG1RF_11099 expression remains unaltered during phage predation of wild type *E. faecalis* OG1RF and Δ*ireK* strains relative to uninfected controls. Data represent three biological replicates. Error bars indicate standard deviation.

**Table S1. List of bacterial strains, phages, plasmids and primers used in this study**.

