## Supplementary material for "Phage infection and sub-lethal antibiotic exposure mediate *Enterococcus faecalis* type VII secretion system dependent inhibition of bystander bacteria": Table S1

**Table S1. List of bacterial strains, phages, plasmids, and primers used in this study**

| **Strains, phages, plasmids, and primers** | **Characteristics and/or description** | **Reference /Source** |
|---|---|---|
| ***Enterococcus faecalis*** | | |
| OG1RF | Human oral isolate; Rf$^R$, Fa$^R$ | [1] |
| $\Delta pip_{V583}$ | V583 background with a deletion of *pip*. Vm$^R$, Em$^R$, Gm$^R$ | [2] |
| Δ*ireK* | *E. faecalis* OG1RF CK119 | [3] |
| Δ*essB* | *E. faecalis* OG1RF markerless deletion in OG1RF_11104 | This study |
| *croR-Tn* | *E. faecalis* OG1RF *croR* transposon mutant. Rf$^R$, Fa$^R$, Cm$^R$ | [4] |
| *croS-Tn* | *E. faecalis* OG1RF *croS* transposon mutant. Rf$^R$, Fa$^R$, Cm$^R$ | [4] |
| *liaR-Tn* | *E. faecalis* OG1RF *liaR* transposon mutant. Rf$^R$, Fa$^R$, Cm$^R$ | [4] |
| *liaS-Tn* | *E. faecalis* OG1RF *liaS* transposon mutant. Rf$^R$, Fa$^R$, Cm$^R$ | [4] |
| OG1RF (pLZ12A) | *E. faecalis* OG1RF carrying pLZ12A empty vector. Rf$^R$, Fa$^R$, Cm$^R$ | This study |
| Δ*essB* (pLZ12A) | Δ*essB* strain carrying pLZ12A empty vector. Rf$^R$, Fa$^R$, Cm$^R$ | This study |
| Δ*essB* (pLZ12A::*essB*) | Δ*essB* strain carrying pLZ12A complementation vector. Rf$^R$, Fa$^R$, Cm$^R$ | This study |
| $\Delta pip_{V583}$ (pLZ12A) | $\Delta pip_{V583}$ strain carrying pLZ12A empty vector. Vm$^R$, Em$^R$, Gm$^R$, Cm$^R$ | [5] |
| $\Delta pip_{V583}$ (pLZ12A::*11110*) | $\Delta pip_{V583}$ strain carrying pLZ12A containing coding sequence of OG1RF_11110 from P-*bacA.* Vm$^R$, Em$^R$, Gm$^R$, Cm$^R$ | This study |
| $\Delta pip_{V583}$ (pLZ12A::*11112*) | $\Delta pip_{V583}$ strain carrying pLZ12A containing coding sequence of OG1RF_11112 from P-*bacA.* Vm$^R$, Em$^R$, Gm$^R$, Cm$^R$ | This study |
| $\Delta pip_{V583}$ (pLZ12A::*11122*) | $\Delta pip_{V583}$ strain carrying pLZ12A containing coding sequence of OG1RF_11122 from P-*bacA.* Vm$^R$, Em$^R$, Gm$^R$, Cm$^R$ | This study |
| $\Delta pip_{V583}$ (pLZ12A::*12413*) | $\Delta pip_{V583}$ strain carrying pLZ12A containing coding sequence of OG1RF_12413 from P-*bacA.* Vm$^R$, Em$^R$, Gm$^R$, Cm$^R$ | This study |
| OG1RF (pCIEtm) | *E. faecalis* OG1RF carrying pCIEtm empty vector. Rf$^R$, Fa$^R$, | This study |

| | | |
|---|---|---|
| | Tc$^R$ | |
| Δ*essB* (pCIEtm) | Δ*essB* strain carrying pCIEtm empty vector. Rf$^R$, Fa$^R$, Tc$^R$ | This study |
| OG1RF_11121-Tn | *E. faecalis* OG1RF *OG1RF-11121* transposon mutant. Rf$^R$, Fa$^R$, Cm$^R$ | [4] |
| OG1RF_11121-Tn (pCIEtm) | OG1RF_11121-Tn strain carrying pCIEtm empty vector. Rf$^R$, Fa$^R$, Cm$^R$, Tc$^R$ | This study |
| OG1RF_11121-Tn (pCIEtm::*11121*) | OG1RF_11121-Tn strain carrying pCIEtm complementation vector. Rf$^R$, Fa$^R$, Cm$^R$, Tc$^R$ | This study |
| Δ*ireK* (pCIE) | Δ*ireK* strain carrying pCIE empty vector. Rf$^R$, Fa$^R$, Cm$^R$ | This study |
| Δ*ireK* (pCIE::*ireK*) | Δ*ireK* strain carrying pCIE complementation vector. Rf$^R$, Fa$^R$, Cm$^R$ | This study |
| OG1RF_11099-Tn | *E. faecalis* OG1RF *OG1RF-11099* transposon mutant. Rf$^R$, Fa$^R$, Cm$^R$ | [4] |
| OG1RF_11099-Tn (pCIEtm) | OG1RF_11099-Tn strain carrying pCIEtm empty vector. Rf$^R$, Fa$^R$, Cm$^R$, Tc$^R$ | This study |
| OG1RF_11099-Tn (pCIEtm::*11099*) | OG1RF_11099-Tn strain carrying pCIEtm complementation vector. Rf$^R$, Fa$^R$, Cm$^R$, Tc$^R$ | This study |
| **Other bacteria** | | |
| *S. aureus* | *Staphylococcus aureus* strain LAC* φ11::LL29 *tet*. Tc$^R$ | [6] |
| *E. faecium* | *Enterococcus faecium* strain 1,231,410. Vm$^R$, Em$^R$ | [7] |
| *L. monocytogenes* | *Listeria monocytogenes* 10403S. St$^R$ | [8] |
| *L. lactis* | *Lactococcus lactis* NZ9000 | Doran lab |
| *S. agalactiae* | *Streptococcus agalactiae* strain COH1. | [9] |
| *S. pyogenes* | *Streptococcus pyogenes* ATCC 12384. | ATCC |
| *S. mitis* | *Streptococcus mitis* NS5. Clinical isolate from UT Southwestern Clinical Microbiology Laboratory | |
| *S. gordonii* | *Streptococcus gordonii* ATCC® 49818. St$^R$ | Doran lab |
| *S. salivarius* | *Streptococcus salivarius* K12. Sp$^R$ | Doran lab |
| *S. enterica* | *Salmonella enterica* serovar Typhimurium. AV09379 *put*::Kan; Kn$^R$ | [10] |
| *V. cholerae* | *Vibrio cholerae C6706 int I4::TnFL63*; Kn$^R$ | [11] |
| ***Escherichia coli*** | | |
| TG1 | *[F' traD36 proAB lacIqZ ΔM15] supE thi-1 Δ(lac-proAB)* | Lucigen |

| | | |
|---|---|---|
| | Δ*(mcrBhsdSM)5(rK - mK -)* | |
| K12 | *Escherichia coli* K12, ATCC 25404 | ATCC |
| **Phage** | | |
| VPE25 | Siphoviridae; Wastewater isolate | [2] |
| **Plasmids** | | |
| pLZ12A | *bacA* promoter cloned into shuttle vector pLZ12; pSH71 origin; $Cm^R$ | [5, 12] |
| pLT06 | *E. faecalis* allelic exchange vector; $Cm^R$ | [13] |
| pBD01 | Δ*essB* construct cloned into pLT06 by Gibson assembly. $Cm^R$ | This study |
| pLZ12A::*essB* | *essB* complementation vector. Cloned into PstI/BamHI site. $Cm^R$ | This study |
| pLZ12A::*11110* | pLZ12A expressing *OG1RF_11110* from $P_{bacA}$. Cloned into PstI/BamHI site. $Cm^R$ | This study |
| pLZ12A::*11112* | pLZ12A expressing *OG1RF_11112* from $P_{bacA}$. Cloned into PstI/BamHI site. $Cm^R$ | This study |
| pLZ12A::*11122* | pLZ12A expressing *OG1RF_11122* from $P_{bacA}$. Cloned into PstI/BamHI site. $Cm^R$ | This study |
| pLZ12A::*12413* | pLZ12A expressing *OG1RF_11122* from $P_{bacA}$. Cloned into PstI/BamHI site. $Cm^R$. | This study |
| pCIE | cCF10 pheromone inducible $P_Q$ expression vector. $Cm^R$ | [14] |
| pCIEtm | Pheromone-inducible pCIE vector with tetracycline resistance cassette. $Tet^R$ | [15] |
| pCIEtm::*11121-11122* | pCIEtm expressing OG1RF_11121-11122 from cCF10 responsive promoter $P_Q$ (11122 is also under control of native promoter). Cloned with BamHI/XbaI into BamHI/NheI sites. $Tet^R$. | This study |
| pCIEtm::*11099* | pCIEtm expressing OG1RF_11099 from cCF10 responsive promoter $P_Q$. Cloned into BamHI/PvuI site. $Tet^R$ | This study |
| pGEM-T-Easy | Cloning vector, $Amp^R$ | Promega |
| pGEM-T-Easy::*ireK* | pGem-T-Easy with *ireK* inserted at the T overhang. $Amp^R$. | This study |
| pCIE::*ireK* | pCIEtm expressing *ireK* from cCF10 responsive promoter | This study |

| | | |
|---|---|---|
| | $P_Q$. Cloned into BamHI/SphI sites. $Cm^R$. | |
| **Primers** | | |
| *essB*-F | NNNNNNCTGCAGATGAGCGATTAAAGGATATTTCA; Forward primer to generate pLZ12A::*essB*; PstI site | This study |
| *essB*-R | NNNNNNGGATCCTTACTATTTTCGTTGTCATCC; Reverse primer to generate pLZ12A::*essB*; BamHI site | This study |
| *OG1RF_11110*-F | NNNNNNCTGCAGATGGACTTCCAAGGTGGTAAAATTAT; Forward primer to generate pLZ12A::*11110*; PstI site | This study |
| *OG1RF_11110*-R | NNNNNNGGATCCTTATTCTCCGTACCATTCCTCTTTA; Reverse primer to generate pLZ12A::*11110*; BamHI site | This study |
| *OG1RF_11112*-F | NNNNNNCTGCAGATGAATAAAATCTTAAATAAAATATCTTTTG; Forward primer to generate pLZ12A::*11112*; PstI site | This study |
| *OG1RF_11112*-R | NNNNNNGGATCCCTAACTATCTTCACCATACCATTCTTG; Reverse primer to generate pLZ12A::*11112*; BamHI site | This study |
| *OG1RF_11122*-F | NNNNNNCTGCAGATGGTTTTCATGATAAAAAATTATGTACC; Forward primer to generate pLZ12A::*11122*; PstI site | This study |
| *OG1RF_11122*-R | NNNNNNGGATCCTTATTTTTTGGTTCTCTTGTTCTTC; Reverse primer to generate pLZ12A::*11122*; BamHI site | This study |
| *11121*-bam-fwd | ATAGGATCCACGTATGTCTAATGAGGAGG; forward primer to amplify OG1RF_11121-11122; BamHI site | This study |
| *11122*-xba-rev | ATATCTAGATTATTTTTTGGTTCTCTTGTTC; reverse primer to amplify OG1RF_11122; XbaI site | This study |
| *ireK*-BamHI-F | GGATCCACCGTGTTAGTGATACA; forward primer to amplify *ireK*; BamHI site | This study |
| *ireK*-SphI-R | GCATGCTTAATTACTCGTACTACT; reverse primer to amplify *ireK*, SphI site | This study |
| *OG1RF_11099*-F | NNNNNNGGATCCATGGTTCAAAATATACCAATTTATATTCAAATTCACG; Forward primer to generate pCIEtm::11099; BamHI site | This study |
| *OG1RF_11099-R* | NNNNNNCGATCGCTACTTCTCTAAATAAAACTCAAATCGACTTCCTGC; Reverse primer to generate pCIEtm::11099; PvuI site | This study |
| *12413*-pst-fwd | ATACTGCAGTAACTATTTTAGGTTCCAGTCC; Forward primer to amplify OG1RF_12413; PstI site | This study |

| | | |
|---|---|---|
| *12413*-bam-rev | TATGGATCCAAAGTATCTGGTATTGTGTTTGC; Reverse primer to amplify OG1RF_12413; BamHI site | This study |
| RT-*esxA*-F | AAGGGCAAGCATTTCAAGCG; qPCR forward primer for OG1RF_11100 | [16] |
| RT-*esxA*-R | TCTTGACGGTCACGTTCTGC; qPCR reverse primer for OG1RF_11100 | [16] |
| RT-*esaA*-F | CCAATGGCTTGGCAACTGAC; qPCR forward primer for OG1RF_11101 | [16] |
| RT-*esaA*-R | GCGAACGAACGTGCATTTTG; qPCR reverse primer for OG1RF_11101 | [16] |
| RT-*essB*-F | GGGAATGGCACCCTGAAAGA; qPCR forward primer for OG1RF_11104 | [16] |
| RT-*essB* -R | CTTCGCGCTTGGCTTTTTGA; qPCR reverse primer for OG1RF_11104 | [16] |
| RT-*essC1*-F | TTGGAAAGGTGGCGGAATAG; qPCR forward primer for OG1RF_11105 | [16] |
| RT-*essC1*-R | TCTGCTTTGATACTGGCTAAGG; qPCR reverse primer for OG1RF_11105 | [16] |
| RT-*11109*-F | GCTTTGGAGAACGCTGAACG; qPCR forward primer for OG1RF_11109 | [16] |
| RT-*11109*-R | TTTTGACAGTCTTGCGCTCG; qPCR reverse primer for OG1RF_11109 | [16] |
| RT-*essC2*-F | CTCAACCGGATCGTGCTTATT; qPCR forward primer for OG1RF_11115 | [16] |
| RT-*essC2*-R | CCTTGGTAGCGAATGGATCATAG; qPCR reverse primer for OG1RF_11115 | [16] |
| RT-*clpX*-F | ATTGGACCAACAGGTTCAGG; qPCR forward primer for *clpX* | This study |
| RT-*clpX*-R | TTTCCGCACGTTCAACATTA; qPCR reverse primer for *clpX* | This study |
| RT-11099-F | GGAACGTATGTAGCACGTAAGA; qPCR forward primer for OG1RF_11099 | This study |
| RT-11099-R | TAAGACACCGTCCGACTAGAA; qPCR reverse primer for OG1RF_11099 | This study |

| | | |
|---|---|---|
| RT-16S-F | CGCTTCTTTCCTCCCGAGT; qPCR forward primer 16S rRNA gene | [16] |
| RT-16S-F | GCCATGCGGCATAAACTG; qPCR reverse primer 16S rRNA gene | [16] |

$Cm^R$ - chloramphenicol resistant; $Rf^R$ - rifampicin resistance; $Fa^R$ - fusidic acid resistance; $Vm^R$ - vancomycin resistance; $Em^R$ - erythromycin resistance; $Gm^R$ - Gentamicin resistance; $Tc^R$ - tetracycline resistance; $St^R$ = streptomycin resistance; $Kn^R$ = Kanamycin resistance; $Sp^R$ = spectinomycin resistance. Restriction enzyme sites are underlined.


1. Bourgogne A, Garsin DA, Qin X, Singh KV, Sillanpaa J, Yerrapragada S, et al. Large scale variation in *Enterococcus faecalis* illustrated by the genome analysis of strain OG1RF. Genome Biol. 2008;9(7):R110. Epub 2008/07/10. doi: 10.1186/gb-2008-9-7-r110. PubMed PMID: 18611278; PubMed Central PMCID: PMCPMC2530867.

2. Duerkop BA, Huo W, Bhardwaj P, Palmer KL, Hooper LV. Molecular basis for lytic bacteriophage resistance in enterococci. MBio. 2016;7(4). Epub 2016/09/01. doi: 10.1128/mBio.01304-16. PubMed PMID: 27578757; PubMed Central PMCID: PMCPMC4999554.

3. Kristich CJ, Wells CL, Dunny GM. A eukaryotic-type Ser/Thr kinase in *Enterococcus faecalis* mediates antimicrobial resistance and intestinal persistence. Proc Natl Acad Sci U S A. 2007;104(9):3508-13. Epub 2007/03/16. doi: 10.1073/pnas.0608742104. PubMed PMID: 17360674; PubMed Central PMCID: PMCPMC1805595.

4. Dale JL, Beckman KB, Willett JLE, Nilson JL, Palani NP, Baller JA, et al. Comprehensive functional analysis of the *Enterococcus faecalis* core genome using an ordered, sequence-defined collection of insertional mutations in strain OG1RF. mSystems. 2018;3(5). Epub 2018/09/19. doi: 10.1128/mSystems.00062-18. PubMed PMID: 30225373; PubMed Central PMCID: PMCPMC6134198.

5. Chatterjee A, Johnson CN, Luong P, Hullahalli K, McBride SW, Schubert AM, et al. Bacteriophage resistance alters antibiotic-mediated intestinal expansion of enterococci. Infect

Immun. 2019;87(6):e00085-19. Epub 2019/04/03. doi: 10.1128/IAI.00085-19. PubMed PMID: 30936157; PubMed Central PMCID: PMCPMC6529655.

11. Cameron DE, Urbach JM, Mekalanos JJ. A defined transposon mutant library and its use in identifying motility genes in *Vibrio cholerae*. Proc Natl Acad Sci U S A.

2008;105(25):8736-41. Epub 2008/06/25. doi: 10.1073/pnas.0803281105. PubMed PMID: 18574146; PubMed Central PMCID: PMCPMC2438431.

12. Perez-Casal J, Caparon MG, Scott JR. Mry, a trans-acting positive regulator of the M protein gene of *Streptococcus pyogenes* with similarity to the receptor proteins of two-component regulatory systems. J Bacteriol. 1991;173(8):2617-24. Epub 1991/04/01. PubMed PMID: 1849511; PubMed Central PMCID: PMCPMC207828.

13. Thurlow LR, Thomas VC, Hancock LE. Capsular polysaccharide production in *Enterococcus faecalis* and contribution of CpsF to capsule serospecificity. J Bacteriol. 2009;191(20):6203-10. Epub 2009/08/18. doi: 10.1128/JB.00592-09. PubMed PMID: 19684130; PubMed Central PMCID: PMCPMC2753019.

14. Weaver KE, Chen Y, Miiller EM, Johnson JN, Dangler AA, Manias DA, et al. Examination of *Enterococcus faecalis* toxin-antitoxin system toxin Fst function utilizing a pheromone-inducible expression vector with tight repression and broad dynamic range. J Bacteriol. 2017;199(12). Epub 2017/03/30. doi: 10.1128/JB.00065-17. PubMed PMID: 28348028; PubMed Central PMCID: PMCPMC5446624.

15. Willett JL, Ji M, Dunny GM. Exploiting biofilm phenotypes for functional characterization of hypothetical genes in *Enterococcus faecalis*. npj Biofilms and Microbiomes volume2019.

16. Chatterjee A, Willett JLE, Nguyen UT, Monogue B, Palmer KL, Dunny GM, et al. Parallel genomics uncover novel enterococcal-bacteriophage interactions. mBio. 2020;11(2). Epub 2020/03/05. doi: 10.1128/mBio.03120-19. PubMed PMID: 32127456; PubMed Central PMCID: PMCPMC7064774.
